# Safety assessment of a novel C-type natriuretic peptide derivative and the mechanism of bone- and cartilage-specific toxicity

**DOI:** 10.1101/655548

**Authors:** Takafumi Yotsumoto, Naomi Morozumi, Ryuichi Nakamura, Toshimasa Jindo, Mayumi Furuya, Yasuyuki Abe, Tomonari Nishimura, Hiroaki Maeda, Hiroyuki Ogasawara, Yoshiharu Minamitake, Kenji Kangawa

## Abstract

ASB20123, a C-type natriuretic peptide/ghrelin chimeric peptide, was designed as a novel peptide and demonstrated full agonistic activity for natriuretic-peptide receptor B and a significantly longer half-life in plasma compared with the native peptide. We researched the toxicological profile of ASB20123, the correlation between the morphological change of the epiphyseal plate and bone and cartilage toxicity, and biomarkers to detect the toxicity. ASB20123 was systemically administered to male and female rats at daily dose levels of 0.5, 1.5, and 5.0 mg/kg/day for 4 weeks. In this study, toxicity was observed as changes related to bone and cartilage tissues, and no other toxicological changes were observed in all animals. Next, ASB20123 was administered to 12-month-old rats with a little epiphyseal plate. The toxic changes related to bone and cartilage tissues were not observed in any animal with a closed epiphyseal plate, indicating that the toxic changes were triggered by the growth-accelerating effect on the bone and cartilage. Furthermore, we searched for the biomarker related to the bone and cartilage toxicity using rats treated with ASB20123 at doses of 0.005, 0.05, 0.5, and 5.0 mg/kg/day for 4 weeks. A close correlation between necrosis/fibrosis in the epiphysis and metaphysis and thickness of the epiphyseal plate in the femur was confirmed in this study. A decrease in the bone mineral density (BMD) of the femur also was associated with the appearance of bone toxicity. These results indicated that the toxicity of ASB20123 was limited to bone- and cartilage-specific changes, and these changes were triggered by an excessive growth accelerating effect. Furthermore, our data suggested that the thickness of the epiphyseal plate and BMD could be reliable biomarkers to predict bone toxicity.

## Introduction

The C-type natriuretic peptide (CNP) analog is one of the most expecting therapeutic approaches to treat achondroplasia [1]. The binding of CNP to natriuretic-peptide receptor B (NPR-B) inhibits fibroblast growth factor receptor 3 downstream signaling [2], recognized as an important regulator of endochondral bone growth [3]. We recently reported that the exogenous administration of CNP-53 has the potential to stimulate skeletal growth related to short stature, and restore the skull morphology and size of foramen magnum in CNP-KO rats [4, 5]. As a CNP derivative, a clinical trial utilizing BMN-111 is currently proceeding in pediatric patients with achondroplasia [6]. We designed ASB20123, a CNP/ghrelin chimeric peptide, as a novel peptide. ASB20123 contains the full-length 22-amino acids of human CNP-22 fused to the 17-amino acids on the C-terminus region of human ghrelin, and the single amino acid is substituted in its ghrelin region. This novel derivative demonstrated full agonistic activity for NPR-B and showed significantly longer half-life in plasma compared with the native forms. A significant and dose-dependent increase in body length was shown in rats after 12 weeks of administration via subcutaneous infusion [7].

CNP is produced in the brain, kidney, bone, blood cells, blood vessels, and heart [8]. NPR-B is expressed in the brain, lung, bone, heart, and ovary tissue. It is also expressed at relatively high levels in fibroblasts and vascular smooth muscle cells [9]. However, the changes that occur after excessive exogenous CNP exposure remain to be clarified. Furthermore, the toxicological profile of the CNP derivative has not been reported previously. In the present study, we evaluated the exhaustive toxicological profile of ASB20123. Furthermore, we researched the relationship between the specific bone and cartilage toxicity and morphological changes of the epiphyseal plate and reliable biomarkers to detect the toxicity.

## Materials and Methods

### Test article

ASB20123 (GLSKGCFGLKLDRIGSMSGLGCVQQRKDSKKPPAKLQPR, C6 and C22 are bound as an intramolecular disulfide bond) was produced by Asubio Pharma Co. Ltd., Japan using a recombinant DNA method in *Escherichia coli*, purified with high-performance liquid chromatography, and verified by amino acid composition analysis and amino acid sequence analysis [7]. The purity of ASB20123 was 98.4%. Acetate buffer (0.03 mol/L) was added with 10 w/v% sucrose and 1 w/v% benzyl alcohol and used as a vehicle. All chemicals and regents used in the present study were purchased from FUJIFILM Wako Pure Chemical Industries, Ltd., Japan and Otsuka Pharmaceutical Factory, Inc., Japan.

### Animals

Sprague-Dawley (SD) rats were purchased from Charles River Laboratories Japan, Inc., Japan and were used for the studies conducted at the Nonclinical Research Center, LSI Medience Corporation, Japan and Asubio Pharma Co., Ltd, Japan. The animals were housed in a humidity- and temperature-controlled environment with an automatic 12-h light/dark cycle. They were provided with a standard, pelleted lab chow diet (CRF-1, Oriental Yeast Co., Ltd., Japan) and tap water *ad libitum*. All animal experiments were conducted in accordance with the Guidelines for Animal Experiments of LSI Medience Corporation, Japan and/or Asubio Pharma Co., Ltd., Japan and were approved by the Institutional Animal Care and Use Committee of LSI Medience Corporation, Japan and/or the Committees for Ethics in Animal Experiments of Asubio Pharma Co., Ltd., Japan, respectively.

### Study protocol and administration

#### Toxicity profiling study (Study 1)

Twenty male and 20 female rats at 7 weeks of age were randomly divided into 4 groups each with 5 males and 5 females. Dose levels were set at 0.5, 1.5, and 5.0 mg/kg/day. The control animals were treated with the vehicle at the same dosing volume as the test article-treated animals. The dosing volume was set at 1 mL/kg, and the dosing volume for each animal was calculated based on the most recent body weight. Each animal was injected with the dosing formulation subcutaneously once a day for 4 weeks into the back of the animals using a disposable injection needle and syringe. The dose levels were selected based on the result of previous study, in which the rats received at the dose of ASB20123 0.15 mg/kg/day for 12 weeks showed over-growth [7].

#### Mechanism study (Study 2)

Fifteen male and 15 female rats at 12 months of age were randomly divided into 3 groups, each with 5 males and 5 females. Dose levels were set at 0.5 and 5.0 mg/kg/day. Other conditions and procedures were the same as those for study 1.

#### Biomarker study (Study 3)

Twenty-five male rats at 7 weeks of age were randomly divided into 5 groups with 5 males each. Dose levels were set at 0.005, 0.05, 0.5, and 5.0 mg/kg/day. Other conditions and procedures were the same as those for study 1.

### Observations and examination items

In study 1, clinical observation, measurements of body weight, food and water consumption, ophthalmology, urinalysis, hematology, blood chemistry, measurements of serum alkaline phosphatase (ALP) isozymes and osteocalcin, body length (naso-anal length), bone mineral density (BMD), organ weight, necropsy, and histopathology analyses were conducted. In studies 2 and 3, clinical observation, measurements of body weight, body length, femur bone length, and BMD, necropsy, and histopathology of the femur and tibia were conducted.

#### Clinical observation

Clinical signs and mortality were observed once or more per day during the administration period.

#### Blood chemistry

Blood samples were collected from the posterior vena cava and centrifuged at 1870 × *g* for 10 minutes to obtain serum samples. The total protein, albumin, A/G ratio, total bilirubin, asparate aminotransferase, alanine aminotransferase, gamma glutamyltranspeptidase, alkaline phosphatase, lactate dehydrogenase, creatine phosphokinase, total cholesterol, triglycerides, phospholipids, glucose, blood urea nitrogen, creatinine, inorganic phosphorus, and calcium were examined with an auto-analyzer (7170, Hitachi Ltd., Japan), and the sodium, potassium, and chloride were examined with an electrolyte analyzer (EA07, A&T Corporation, Japan).

#### ALP isozymes and osteocalcin

The remaining serum samples collected for blood chemistry were used for the serum ALP isozyme and osteocalcin measurement. For ALP isozyme measurement, the auto electrophoresis system (Epalyzer 2, Helena Laboratories Co., Ltd., USA) was used. For osteocalcin measurement, an immunoradiometric assay was applied with the Rat Osteocalcin IRMA kit (Immutopics, International, LLC., USA).

#### Body length (naso-anal length and femur)

At the end of the administration period, naso-anal lengths of each rat were examined with a scale after euthanasia. Femoral lengths were measured with digital calipers after removal at necropsy.

#### Bone mineral density (BMD)

Bone mineral density (cortical and sponge) of the isolated left femur of each rat was measured using CT scanning (Latheta LCT-200; Hitachi Aloka Medical, Japan).

#### Histological examination

Whole rats specimens were embedded in paraffin, sectioned, stained with hematoxylin and eosin (HE), and examined microscopically. The thickness of the epiphyseal plate at the proximal end of the femur was measured under a light microscope. It was measured at nine sites for the proximal end of the femur. The average thickness was considered the epiphyseal plate thickness for each rat.

### Statistical Analysis

The homogeneity of variance was tested by Bartlett’s method. When the groups were accepted as homogeneous, Dunnett’s multiple comparison test was used for comparison of groups of data. When the groups of data were shown to be heterogeneous, Steel’s multiple comparison test was applied to mean values. A two-tailed test was used as Bartlett’s test, and *P* values less than 0.05 were considered statistically significant.

## Results

### Toxicity profile of ASB20123 in study 1

In the clinical observation, abnormal gait appearing as a shuffling gait of the hind limbs was observed in all the test article-treated groups at the dose level of 0.5 mg/kg/day or more at the 3^rd^ week of administration and later. The number of animals exhibiting these symptoms increased dose-dependently. The BMD values of both cortical and spongy bone in the femur were significantly low in all the test article-treated groups compared with the control group (Fig 1).

**Fig 1.**
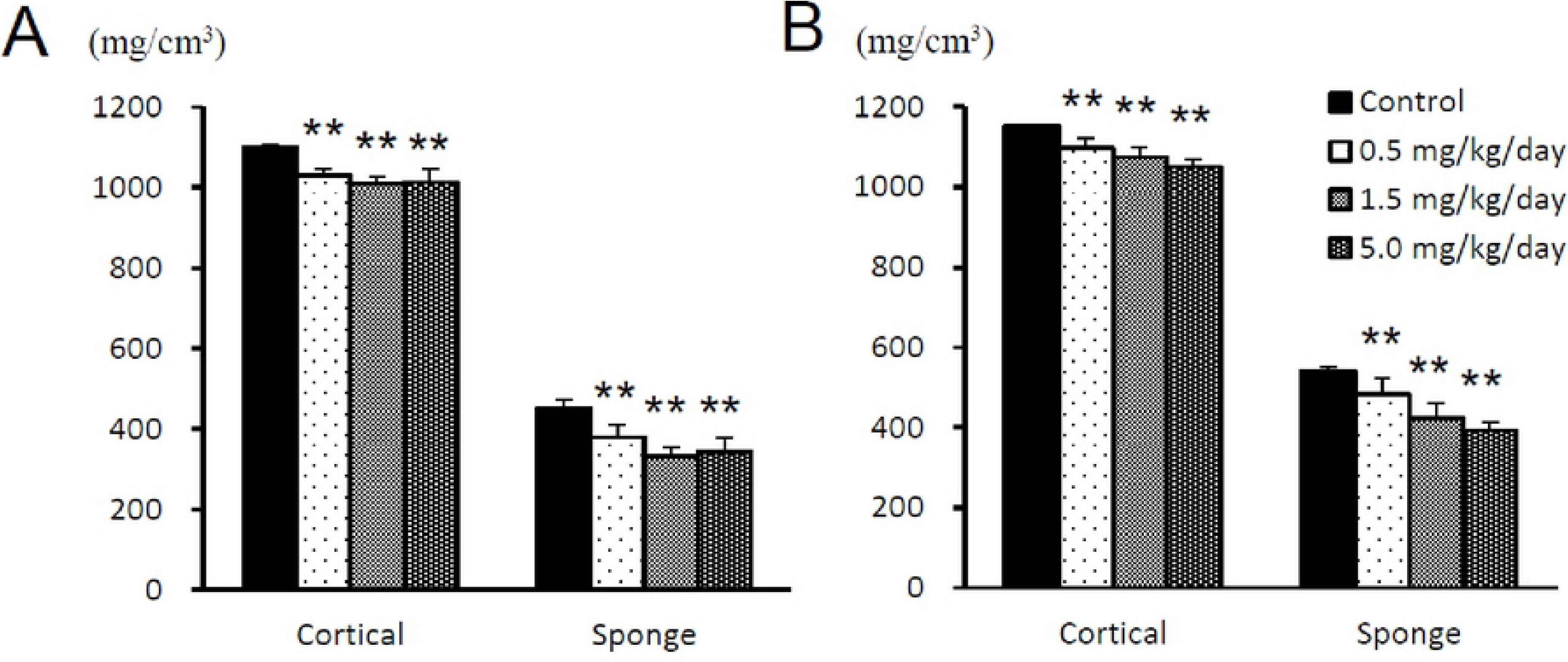
BMD of both the cortical and spongy bone in the femurs of male (A) and female (B) rats treated subcutaneously with ASB20123 for 4 weeks. Each value represents the mean ± SD of 5 rats, ** *P* < 0.01 vs. vehicle-treated group by Dunnett’s multiple comparison test.

The results of histopathological examination are shown in S1 table, and representative histopathological findings for the proximal femoral bone in rats are shown in Fig 2. The test article-related changes were observed in the bone/cartilage tissues as follows. In the femur, thickening of the epiphyseal plate was observed at the dose levels of 0.5 mg/kg/day or more. This change involved both the proximal and distal portions of the femur, was intense in the proximal portion, and was concomitant with increases in the primary bone and osteoblasts. In the proximal portion, there was necrosis of the epiphysis/metaphysis, fibrosis in the marrow of the head, and ectopic chondrogenesis/osteogenesis. In the tibia, thickening of the epiphyseal plate and an increase in the primary bone were observed in all the test article-treated groups. These changes involved both the proximal and distal portions of the tibia and were intense or highly frequent in the distal portion. There were also increases in osteoblasts in the proximal and distal portions in males, and necrosis of the epiphysis/metaphysis and inflammatory changes in the surrounding tissues in the distal portion.

**Fig 2.**
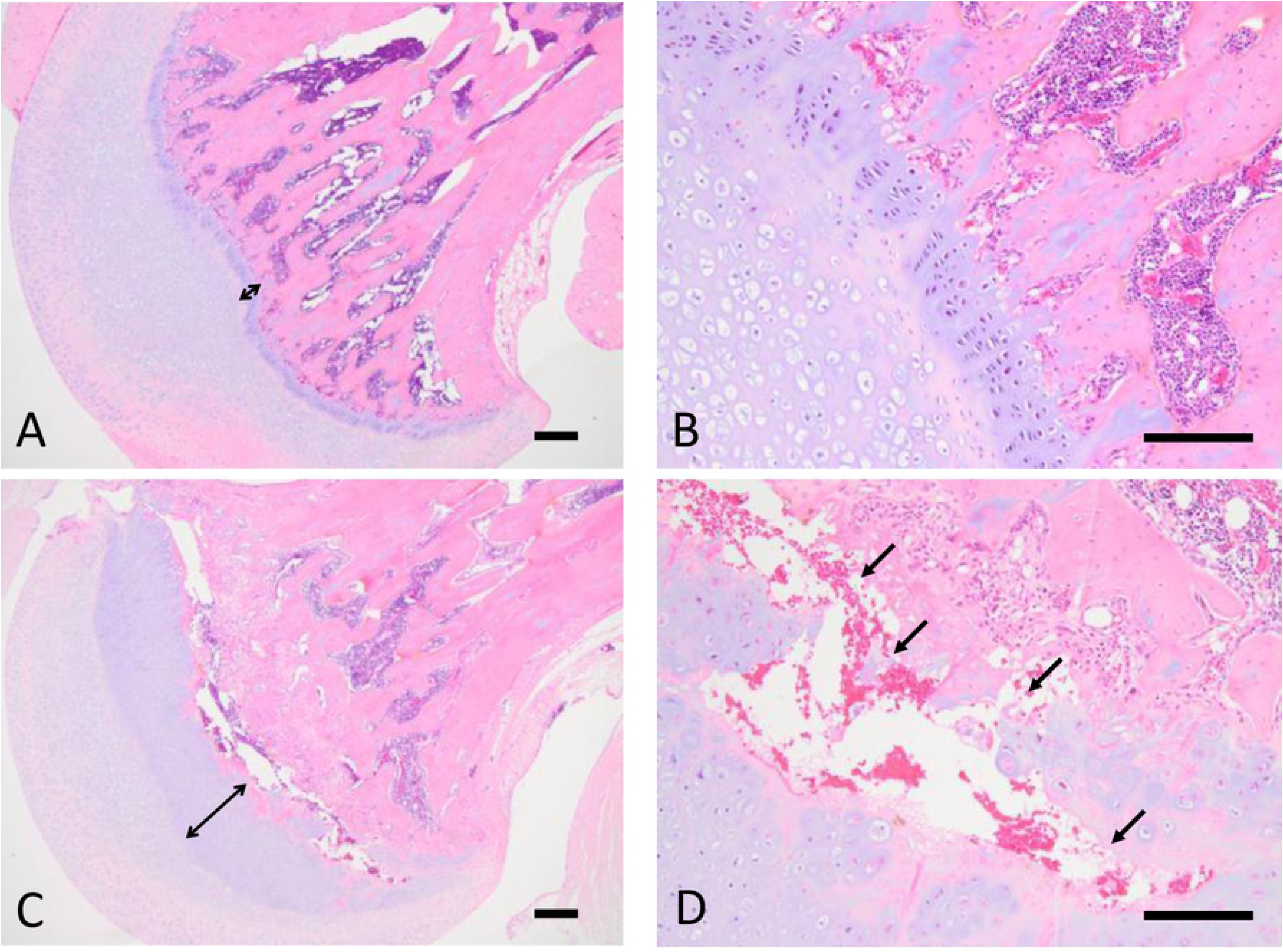
Representative histopathological findings in the proximal femoral bone in rats. (A) Vehicle group (× 40). (B) Vehicle group (× 100). (C) 0.5 mg/kg/day group (× 40). (D) 0.5 mg/kg/day group (× 100). Bidirectional arrows indicate the width of the epiphyseal plate. Arrows indicate the necrosis of cartilage/osseous tissues. Scale bars represent 200 μm.

The changes in body length, ALP activity, ALP-isozyme fraction, and osteocalcin values are shown in Fig 3. Significantly high values of body length were shown in males at the dose levels of 5.0 mg/kg/day and in females at 0.5 mg/kg/day and more, compared with the vehicle control group. The ALP activity was significantly high in males at the dose levels of 1.5 and 5.0 mg/kg/day. Although there was no statistically significant difference, the same tendency was observed in males at the dose levels of 0.5 mg/kg/day and in females at 1.5 and 5.0 mg/kg/day. The ALP-isozyme fraction 3 was significantly high in males and females at the dose levels of 5.0 mg/kg/day compared with the vehicle control group, and the same tendency was observed in males administered 0.5 and 1.5 mg/kg/day and females administered 1.5 mg/kg/day. Meanwhile, there were no changes in the osteocalcin concentrations in any group compared to the control group.

**Fig 3.**
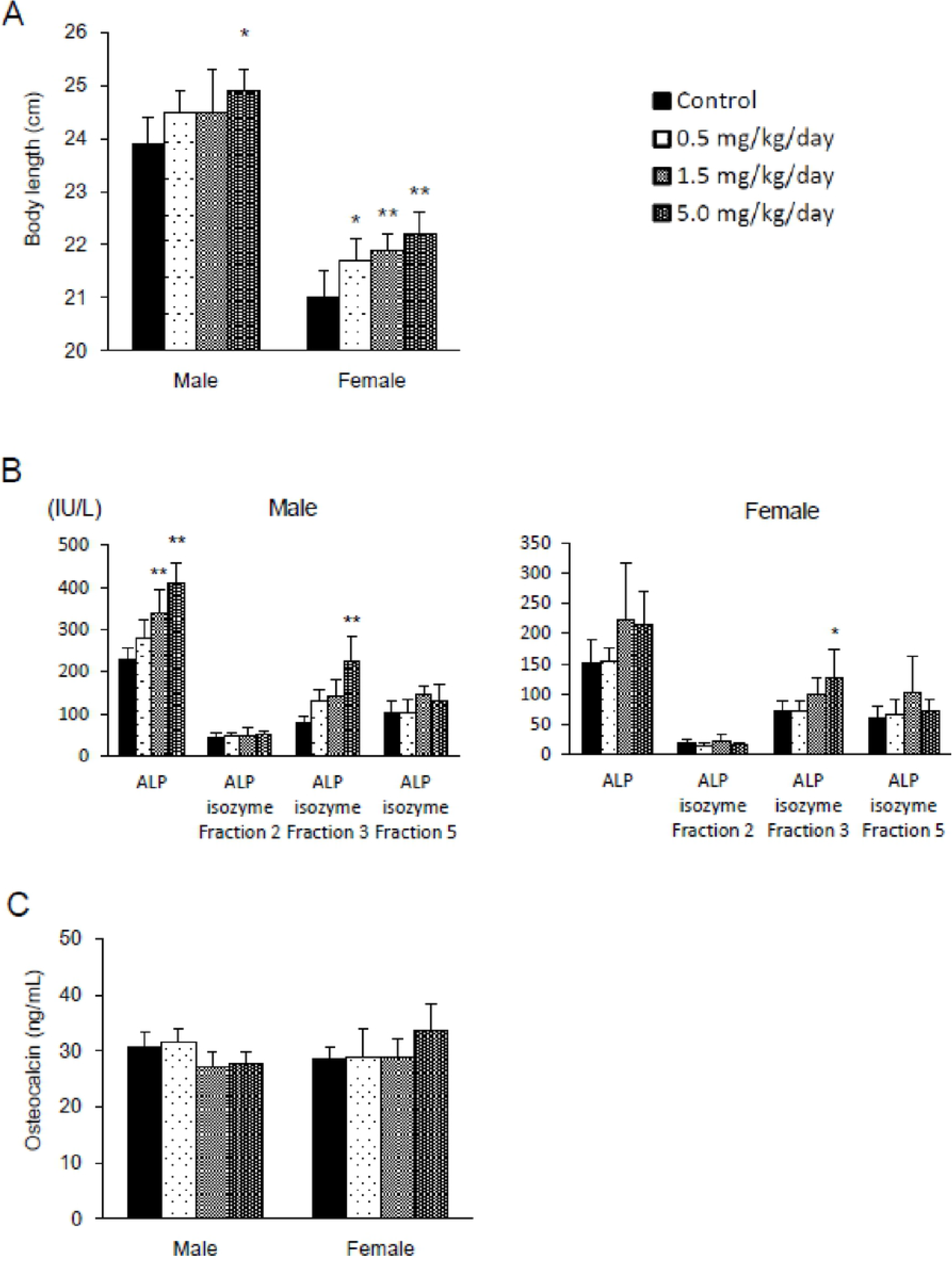
Effects of ASB20123 on the body length (A), ALP and ALP-isozyme fraction activity (B), and osteocalcin value (C) in rats treated subcutaneously for 4 weeks. Each value represents the mean ± SD of 5 rats, * *P* < 0.05, ** *P* < 0.01 vs. vehicle-treated group by Dunnett’s multiple comparison test.

No changes were observed in any group in body weight, food and water consumption, ophthalmology, urinalysis, hematology, necropsy, and organ weight.

### Mechanism of the specific bone and cartilage toxicity in study 2

ASB20123 was administered to the 12-month-old rats. The epiphyseal plate closure was observed in all observation sites of the vehicle group of the aged rats, except for the proximal tibia in 2 female rats. The number of rats with the remaining epiphyseal plate in the treatment group was larger than that in the vehicle group. Increases in osteoblasts and primary bone and degeneration/necrosis in the epiphysis and metaphysis were observed in some animals with the epiphyseal plate, but these findings were not observed in animals with a closed epiphyseal plate (Table 1). No test article-related changes were observed in the clinical observation, body weight, femur bone length, and BMD, but abnormal gait was observed in only 1 animal in the clinical observation in the 0.5 mg/kg/day dosage group.

**Table 1.**
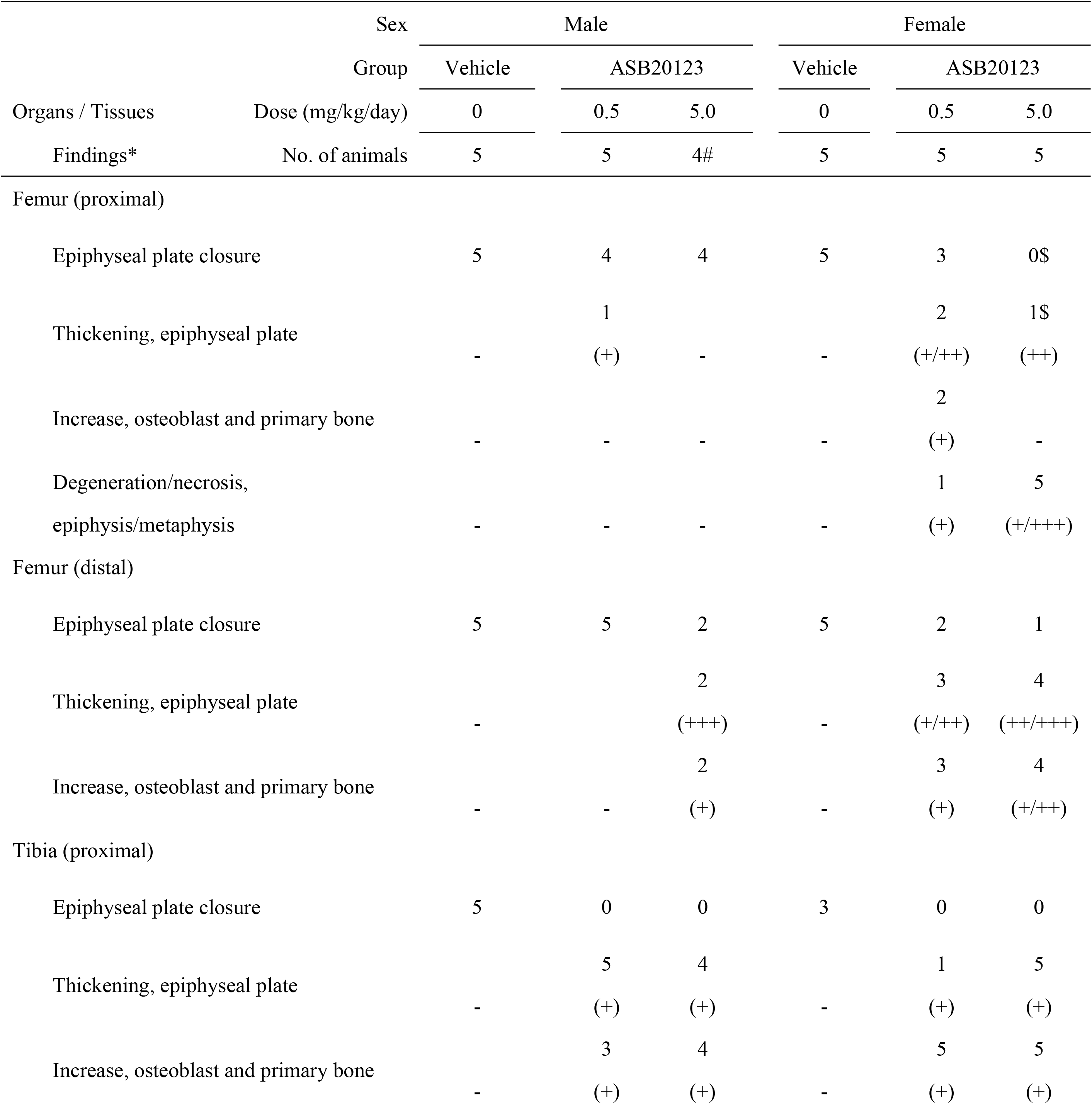
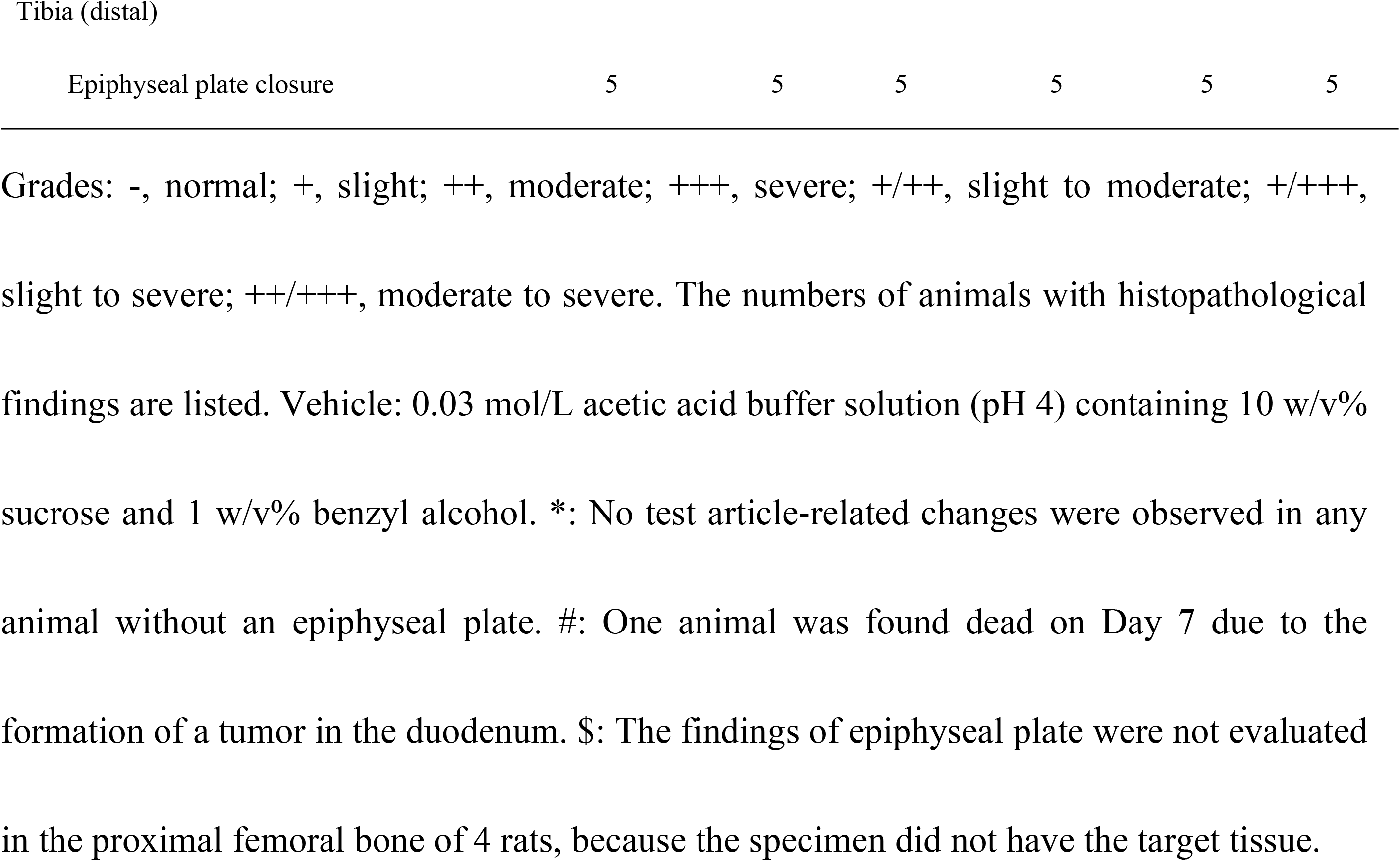
Histopathological findings in the femur and tibia in study 2

**Table 2.**
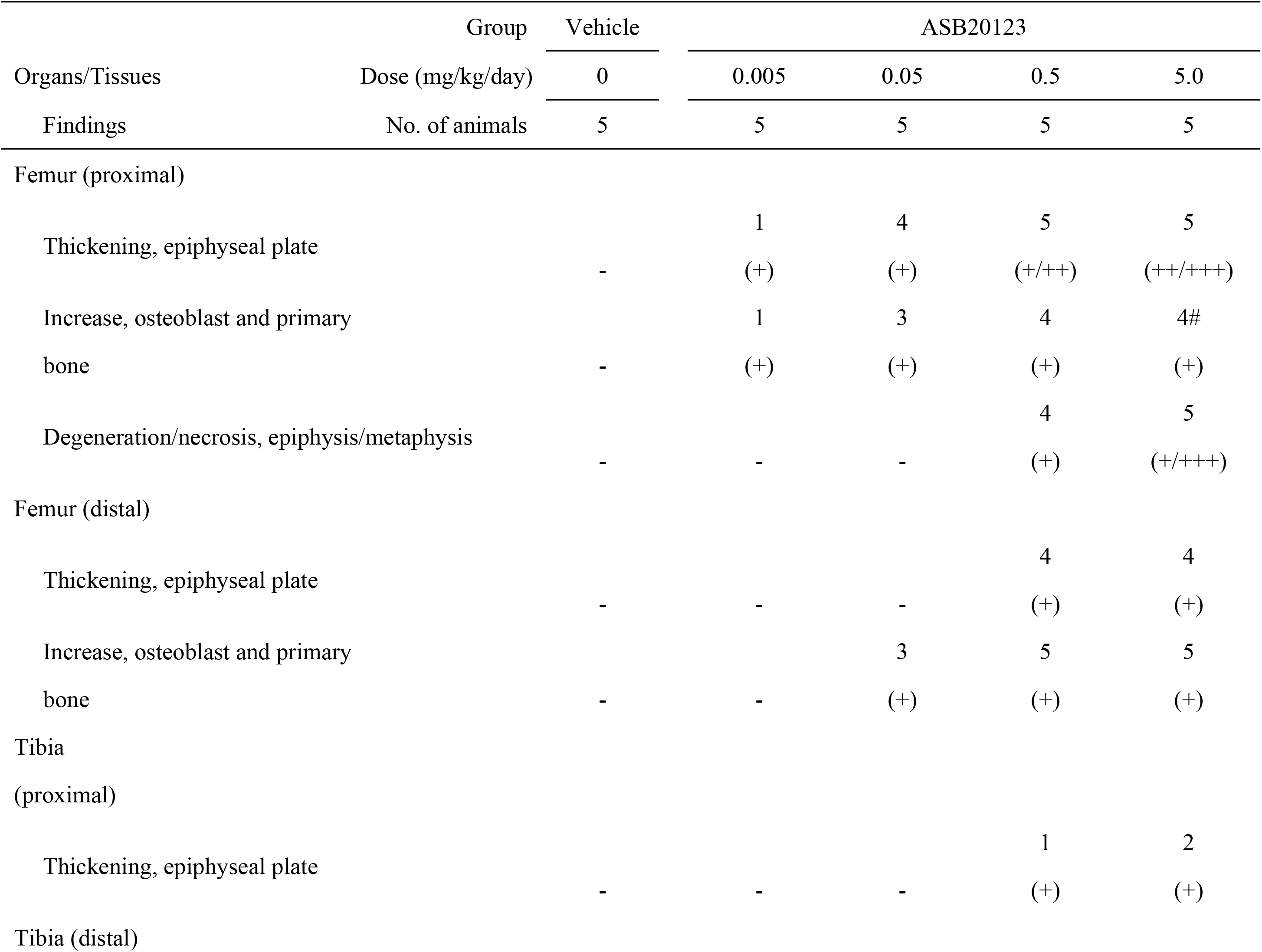
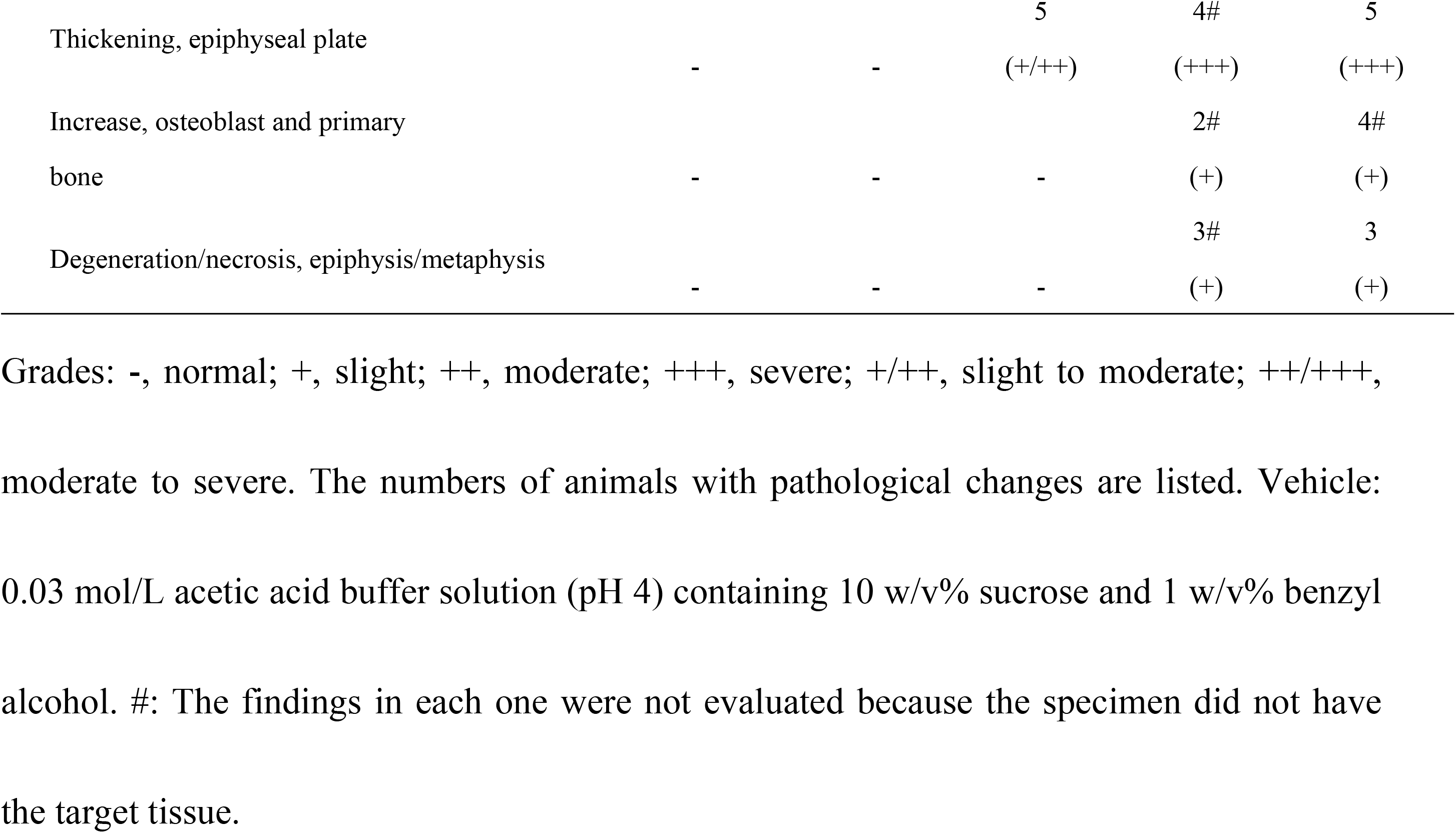
Histopathological findings of the femur and tibia in study 3

### Searching for the biomarker related to the bone and cartilage toxicity in study 3

The thickness of the epiphyseal plate, femur bone length, and bone mineral density of the femur were measured in the rats treated with ASB20123 at doses of 0.005, 0.05, 0.5, and 5.0 mg/kg/day for 4 weeks. The epiphyseal plate thickness increased in a dose-dependent manner, and the toxic findings in the epiphysis and metaphysis were observed only in individuals with thickening of more than 200 μm of the epiphyseal plate. A decrease in the BMD in the femur also reflected the appearance of bone toxicity. In contrast, a correlation between the appearance of bone toxicity and body or femur lengths was not observed (Fig 4). The results for the other observations and examination items were similar to those of study 1.

**Fig 4.**
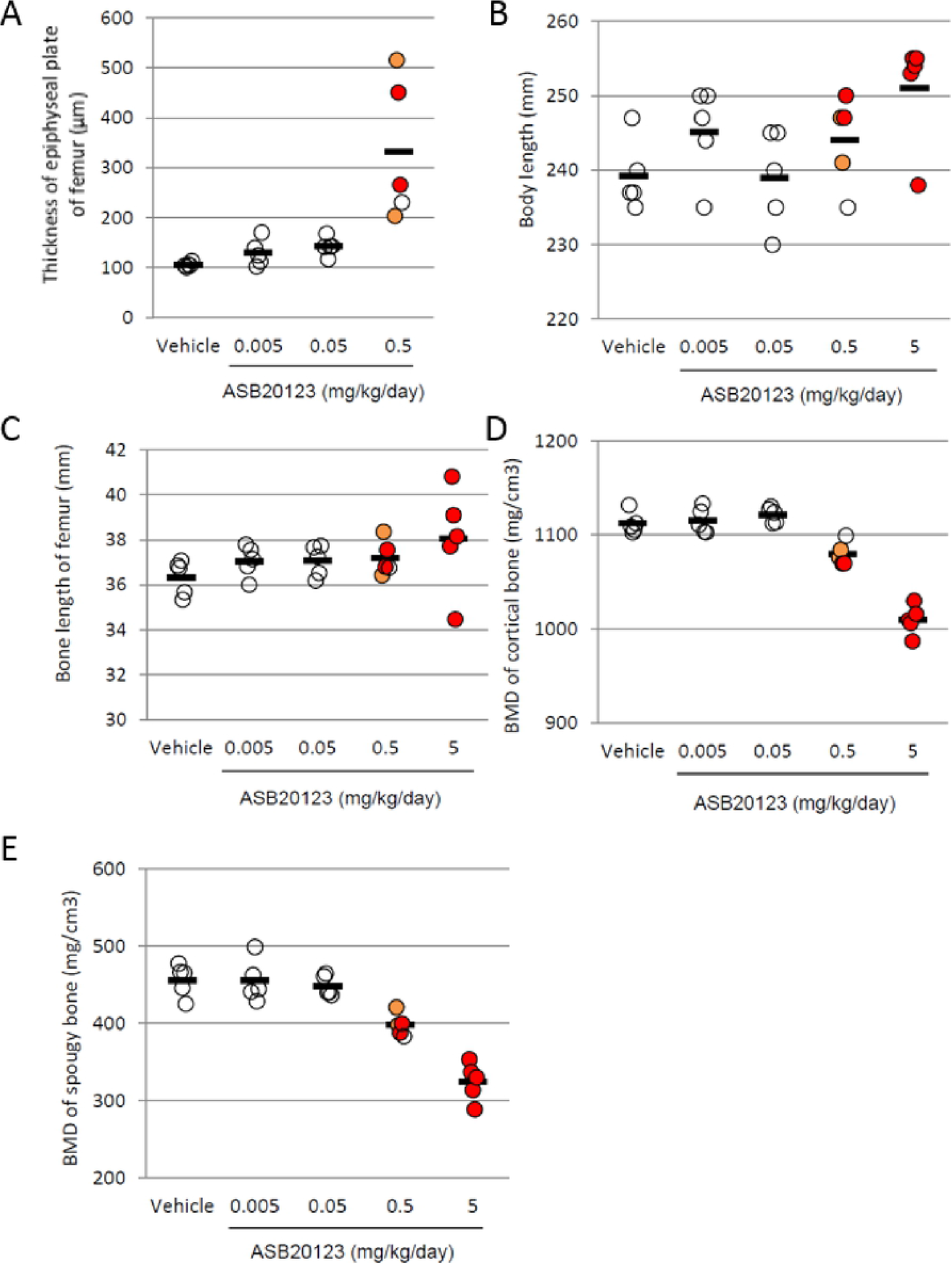
The correlation between bone and cartilage toxicity and several parameters. The thickness of the epiphyseal plate of the femur (A), body length (B), and bone length of the femur (C). The BMD of the cortical bone (D) and spongy bone (E) in the femurs is shown. Bone toxicity observed in each animal is shown in the colored circle, the open circle represents no toxicity, orange indicates slight toxicity, and red indicates severe toxicity. Each bar represents the mean of 5 rats.

## Discussion

ASB20123 is a CNP derivative, and this peptide stimulates bone growth through proliferation and differentiation of chondrocytes [7]. In this study, ASB20123 was administered to male and female rats at daily dose levels of 0.5, 1.5, and 5.0 mg/kg/day for 4 weeks to investigate its toxicity. In this study, toxic changes were observed in the bone and cartilage tissues, and no other toxic changes were observed in all animals.

In the histopathological examination, thickening of the epiphyseal plate, which was frequently accompanied by increases in the primary bone and osteoblasts, was observed in the femur and tibia in all the test article-treated groups. Similar cartilage thickening was also detected in the temporomandibular joint and sternum. These findings were characteristic, prominent, and therefore considered to be primary changes based on the pharmacological action of ASB20123. In relation to the above osseous changes, the body length was extended in all the test-article treated groups, and serum ALP-isozyme fraction activity increased, since it is derived from bone and is elevated in the serum as a result of various bone diseases and bone growth. In addition to the above changes due to a pharmacological action of ASB20123, the following toxicity findings were observed in this study. In the proximal portion of the femur, there was necrosis of the epiphysis/metaphysis, fibrosis in the marrow of the head, and ectopic chondrogenesis/osteogenesis in all the test article-treated groups. Necrosis of the trabecula/marrow in the epiphysis of the head and degeneration/necrosis of the peripheral muscle fibers was also sporadically observed. In the tibia, there was necrosis of the epiphysis/metaphysis and inflammatory changes in the surrounding tissues in the distal portion. All these findings associated with the above-mentioned primary changes were probably inflammatory, ischemic, or reactive due to the physical or physiological stimulation following thickening of the epiphyseal plate. These changes were intense or localized in the proximal portion, especially in the head of the femur and in the distal portion of the tibia, suggesting a possibility that the landing-shock on the hind limbs during walking/moving accelerates these bone/cartilage changes. Clinical signs revealed shuffling gait of the hind limbs. This symptom was considered to be caused by the excessive cartilage increase in the distal tibia.

To research the involvement of the epiphyseal plate in the bone-related changes, ASB20123 was administered to the 12-month-old rats with a little epiphyseal plate. The epiphyseal plate closure was observed in all examination sites of the vehicle group, except for in the proximal tibias of 2 female rats. The number of rats with the remaining epiphyseal plate in the test article-treated group was larger than that in the vehicle group. It was suggested that the administration of ASB20123 delayed the epiphyseal plate closure, and this result corresponded to those of our previous reports [4, 5]. An increase in osteoblasts and primary bone and degeneration/necrosis in the epiphysis and metaphysis were not observed in any animal with a closed epiphyseal plate. These results indicated that the toxic changes in the bone and cartilage tissues were triggered by the excessive growth-accelerating effect based on the pharmacological action of ASB20123.

Biomarkers related to bone and cartilage toxicity, thickness of the epiphyseal plate, body length, femur bone length, and BMD of the cortical and spongy bone in the femur were measured in the rats treated with ASB20123 at the doses of 0.005, 0.05, 0.5, and 5.0 mg/kg/day for 4 weeks. A reliable correlation between necrosis/fibrosis in the epiphysis and metaphysis and thickness of the epiphyseal plate of the femur was confirmed in this study. A decrease in BMD in the cortical bone in the femur was also relevant to the bone and cartilage toxicity. These parameters might be good markers to predict bone- and cartilage-specific toxic changes, because the thickness of the epiphyseal plate can be monitored using radiographic examination, computed tomography, and magnetic resonance imaging in humans [12, 13].

In this study, we evaluated the toxic profile of ASB20123 the CNP derivative with an extended half-life. As a result, over-dosing of ASB20123 induced excessive growth acceleration through endochondral bone growth, resulting in bone- and cartilage-specific toxicity changes in normal young rats without closed epiphyseal plates. Furthermore, our data suggested that the thickness of the epiphyseal plate and BMD of the cortical bone could be reliable biomarkers to predict bone- and cartilage-specific toxicity. The dosage regimen of CNP analog and derivative would be a key factor for successful of therapeutic drug.

## Supporting Information

S1 Table Histopathology in rats treated subcutaneously with ASB20123 for 4 weeks.

